# Solute Carrier Transporter Family Modulates Neutrophil Metabolism During Health and Disease

**DOI:** 10.64898/2026.05.20.726273

**Authors:** Sunayana Malla, Rajib Saha

## Abstract

Neutrophils are the most abundant leukocytes in humans and play a central role in immune regulation. Although traditionally viewed as terminally differentiated cells with limited plasticity, growing evidence indicates that neutrophils exhibit substantial functional heterogeneity in response to stress. To date, however, most studies have focused on transcriptional and signaling changes, while metabolic heterogeneity, especially beyond central carbon metabolism, remains poorly characterized. Here, we systematically investigate metabolic reprogramming in neutrophils under three stress conditions: granulocyte colony-stimulating factor (G-CSF) treatment, hematopoietic stem cell transplantation (HSCT), and pancreatic ductal adenocarcinoma (PDAC). Using condition-specific genome-scale metabolic (GSM) models, we identify distinct metabolic vulnerabilities across neutrophil states. Vitamin metabolism emerged as a key differentiating feature between G-CSF- and HSCT-treated neutrophils, whereas PDAC-associated neutrophils displayed globally enhanced metabolic activity coupled with restricted metabolite exchange fluxes. Furthermore, solute carrier (SLC) family transporters were identified as major metabolic regulators underlying stress-induced neutrophil reprogramming. Together, our findings demonstrate that neutrophil heterogeneity extends beyond transcriptional programs to encompass profound metabolic specialization, highlighting metabolism as a critical dimension of neutrophil plasticity in health and disease.

## Introduction

Neutrophils are the most abundant leukocytes in the human body and, despite their short lifespan, are continuously produced to ensure effective host defense against infection, tissue injury, and inflammation^1^. Neutrophils are generated in the bone marrow through the tightly controlled process of granulopoiesis and exert their antimicrobial activity through mechanisms such as phagocytosis, oxidative burst marked by heightened production of reactive oxygen species (ROS), and the formation of neutrophil extracellular traps (NETs)^1,2^. They constitute the first line of the innate immune response acting as the crucial, and immediate response before adaptive immune system fully activates^3^. Over the past decade, the long-standing view of neutrophils as simple, short-lived effector cells has been fundamentally revised^4,5^. Accumulating evidence demonstrates that neutrophils possess a wide range of functions and engage in extensive crosstalk with other immune cells^2,6^. They are now recognized as highly complex and heterogeneous, exhibiting diverse phenotypic and functional states, particularly in pathological settings such as inflammation and cancer, where they can actively contribute to disease progression^2,5,7^. Despite these advances, the contribution of cellular metabolism to the regulation and diversification of neutrophil functions remains relatively underexplored^8^.

Neutrophils are known to dynamically reprogram their metabolic pathways to support distinct effector functions, including chemotaxis, ROS generation, NET formation, and degranulation^4,9^. Core metabolic pathways that have been studied in this context include glycolysis/gluconeogenesis, the pentose phosphate pathway (PPP), fatty acid synthesis and oxidation (FAS/FAO), glutaminolysis, and the tricarboxylic acid (TCA) cycle coupled with oxidative phosphorylation (OXPHOS)^4,9,10^. Shifts within these major pathways play a critical role in shaping neutrophil behavior during disease initiation, resolution, and progression^9^.

However, direct comparisons of metabolic reprogramming across neutrophils under different critical conditions such as following Granulocyte Colony-Stimulating Factor (G-CSF) treatment, hematopoietic stem cell transplantation (HSCT), or in disease contexts like pancreatic ductal adenocarcinoma (PDAC) remain limited. Understanding the metabolic heterogeneity of neutrophils across these settings is essential, given their central role in regulating neutrophil development and function in both health and disease^11^. For instance, granulocyte colony-stimulating factor (G-CSF) is extensively utilized to stimulate neutrophil production in conditions such as neutropenia^12,13^. On the other hand, hematopoietic stem cell transplantation (HSCT) is employed in diseases like multiple sclerosis (MS) to restore a dysregulated immune system^14,15^. Although both treatments are highly effective, the neutrophils generated afterward may exhibit distinct functional properties^16,17^. Furthermore, achieving a comprehensive understanding of neutrophil immunometabolism necessitates expanding beyond central carbon metabolism to encompass secondary and auxiliary metabolic pathways^4^.

In this regard, systems biology approaches, particularly genome-scale metabolic (GSM) modeling, provide a powerful framework for achieving a holistic view of neutrophil metabolism^18,19^. These models integrate biochemical reactions, metabolites, and genes to elucidate how metabolic networks collectively drive specific cellular phenotypes^20^. GSM models are comprehensive mathematical representations of an organism’s metabolic network, linking genes to enzymatic reactions and metabolites through gene–protein–reaction (GPR) associations^21^. By incorporating context-specific experimental data such as transcriptomics, metabolomics, and single-cell RNA sequencing, we can ensure the construction of biologically accurate and predictive models^19^. Over time, these models have become indispensable tools for investigating metabolic adaptations across diverse biological systems, ranging from simple microorganisms to complex human immune cells^22–25^, while also offering valuable mechanistic insights into disease progression. Although significant advances have been made in developing metabolic models for immune cells such as macrophages and mast cells leading to important discoveries, similar efforts focused on neutrophils remain comparatively scarce^8^.

In this study, we reconstructed four genome-scale metabolic (GSM) models of neutrophils under Normal, Granulocyte colony-stimulating factor (G-CSF)–treated, Hematopoietic stem cell transplantation (HSCT)-treated, and Pancreatic Cancer (PDAC) conditions by integrating available single-cell RNA-sequencing data^8^. To ensure biological fidelity, each reaction in the models was constrained using condition-specific bounds derived from GPR associations. The resulting models successfully captured key metabolic features of neutrophils across conditions, including enhanced carbohydrate metabolism in both G-CSF– and HSCT-treated neutrophils, as well as condition-specific alterations in tricarboxylic acid (TCA) cycle reactions in the PDAC model. Beyond primary metabolism, we performed an extensive comparative analysis of multiple metabolic pathways, revealing condition-dependent metabolic rewiring relative to normal neutrophils. Notably, our analysis identifies Solute Carrier (SLC) family transporters as critical modulators of global neutrophil metabolic activity in response to diverse stressors, including G-CSF stimulation, HSCT treatment, and Pancreatic Cancer. While these GSM models rely on pseudo–steady-state assumptions and therefore cannot capture the full temporal complexity of immune system dynamics, they nevertheless highlight the substantial metabolic heterogeneity exhibited by neutrophils under different stress conditions and underscore the central role of SLC transporters in regulating neutrophil metabolism.

## Results

### Generation and Validation of Condition-Specific Neutrophil Metabolic Models

Four condition-specific neutrophil metabolic models were reconstructed from the global human metabolic network, Human1^26^, by integrating scRNA-seq data corresponding to Normal, G-CSF–treated, HSCT-treated, and Pancreatic Cancer (PC) conditions using the ftINIT pipeline^8,27^. Additionally, gene expression (obtained from Montaldo et al., 2022) were incorporated to constrain reaction bounds following model generation, resulting in metabolically distinct models for each condition as shown in **Figure 1a**. This integration enabled the construction of biologically responsive models capable of capturing condition-specific metabolic behavior such as Reactive Oxygen Species (ROS) production, Fatty acid metabolism, and Tricarboxylic Acid (TCA) cycle. The details of each metabolic model are available in **the Supplementary File**.

**Figure 1:**
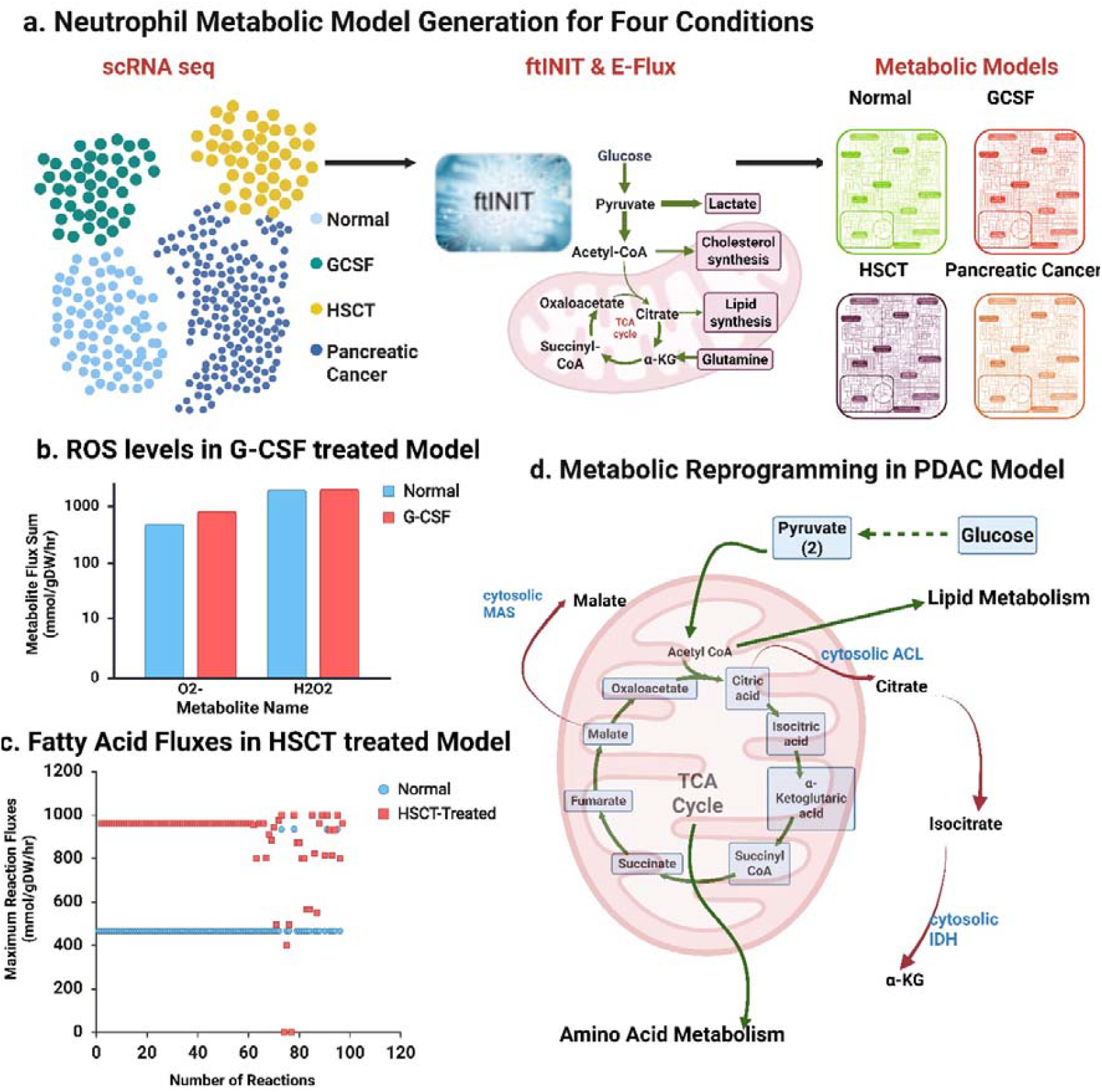
Generation and validation of condition-specific neutrophil metabolic models. Single-cell RNA-sequencing data for neutrophils under four conditions, Normal, G-CSF–treated, HSCT-treated, and Pancreatic Cancer (PDAC), were obtained from Montaldo et al. (2022). The ftINIT pipeline was used to construct four genome-scale metabolic models, with each reaction constrained by condition-specific bounds derived from gene–protein–reaction (GPR) associations. This approach enabled the generation of biologically sensitive metabolic models. The resulting models were validated by their ability to recapitulate known neutrophil biology under each condition. (b) G-CSF treated neutrophils showed a slight increase in H2O2, AND O2-pool as reported in literature. (c) As reported by Machado et al., 2022, the HSCT treated neutrophils showed enhanced fluxes in reactions from fatty acid metabolism. (d) Neutrophils from the pancreatic cancer condition displayed heightened overall metabolic activity, particularly in mitochondrial pathways, accompanied by reduced flux through specific reactions, including the transport of malate, citrate, and α-ketoglutarate. The green arrows show the increased flux space, and red arrows show reduced flux space in PDAC neutrophils in comparison to normal condition.

First, we evaluated whether the reconstructed neutrophil models could recapitulate known biological phenomena (as mentioned above) associated with each condition. To this end, we compared changes in reaction flux ranges obtained from Flux Variability Analysis (FVA)^28,29^, using the Normal condition as the baseline. As reported by Montaldo et al., 2022 we found increased O^-^_2_ and H_2_O_2_ in G-CSF treated neutrophils (**Figure 1b**), increased fatty acid activation (cytosolic) in HSCT treated neutrophils (**Figure 1c**)^8^. Additionally, we also observed increased flux space in reactions from carbohydrate metabolism (such as glycolysis/gluconeogenesis, Starch and sucrose metabolism, fructose and mannose metabolism, etc), in both G-CSF–treated and HSCT-treated neutrophils^30^. In the PDAC neutrophil models, we observed expanded flux ranges in glycolysis, and lipid metabolism along with core mitochondrial TCA cycle reactions, alongside reduced flux capacity in cytosolic ATP-citrate lyase, cytosolic IDH1, and the cytosolic malate dehydrogenase shuttle (**Figure 1d**)^25,31^. These patterns closely reflect the metabolic phenotype of neutrophils in cancer, which is characterized by a highly active, high-flux TCA cycle coupled with oxidative phosphorylation, and minimal to near-absent activity in cytosolic lipogenic pathways. Given that neutrophils are intrinsically non-lipogenic, the suppression of ACLY, IDH1, and malate dehydrogenase–mediated cytosolic fluxes is expected as shown in **Figure 1d**^32,33^.

Collectively, the ability of the models to reproduce these established metabolic states provided confidence to further interrogate neutrophil metabolic reprogramming across conditions and to identify key regulators governing neutrophil metabolism. The details of the flux comparison are available in **the Supplementary File**.

### Metabolic Reprogramming of Neutrophil in G-CSF treated and HSCT treated Neutrophils

Our flux analysis provides insight into the metabolic reprogramming of G-CSF–treated and HSCT-treated neutrophils. We observed that G-CSF–treated cells display significantly different metabolic behaviors compared with neutrophils under normal conditions and, notably, show greater metabolic similarity to neutrophils from HSCT-treated conditions (**Figure 2a)**. Specifically, we identified expanded flux spaces in reactions associated with pathways such as acyl-CoA hydrolysis and bile acid biosynthesis, fatty acid activation (cytosolic) along with increased exchange of di- and tripeptides. However, approximately 70% of reactions in the model exhibited a reduced flux space compared with the normal condition, indicating a broad contraction of metabolic flexibility despite localized pathway expansions as shown.

**Figure 2:**
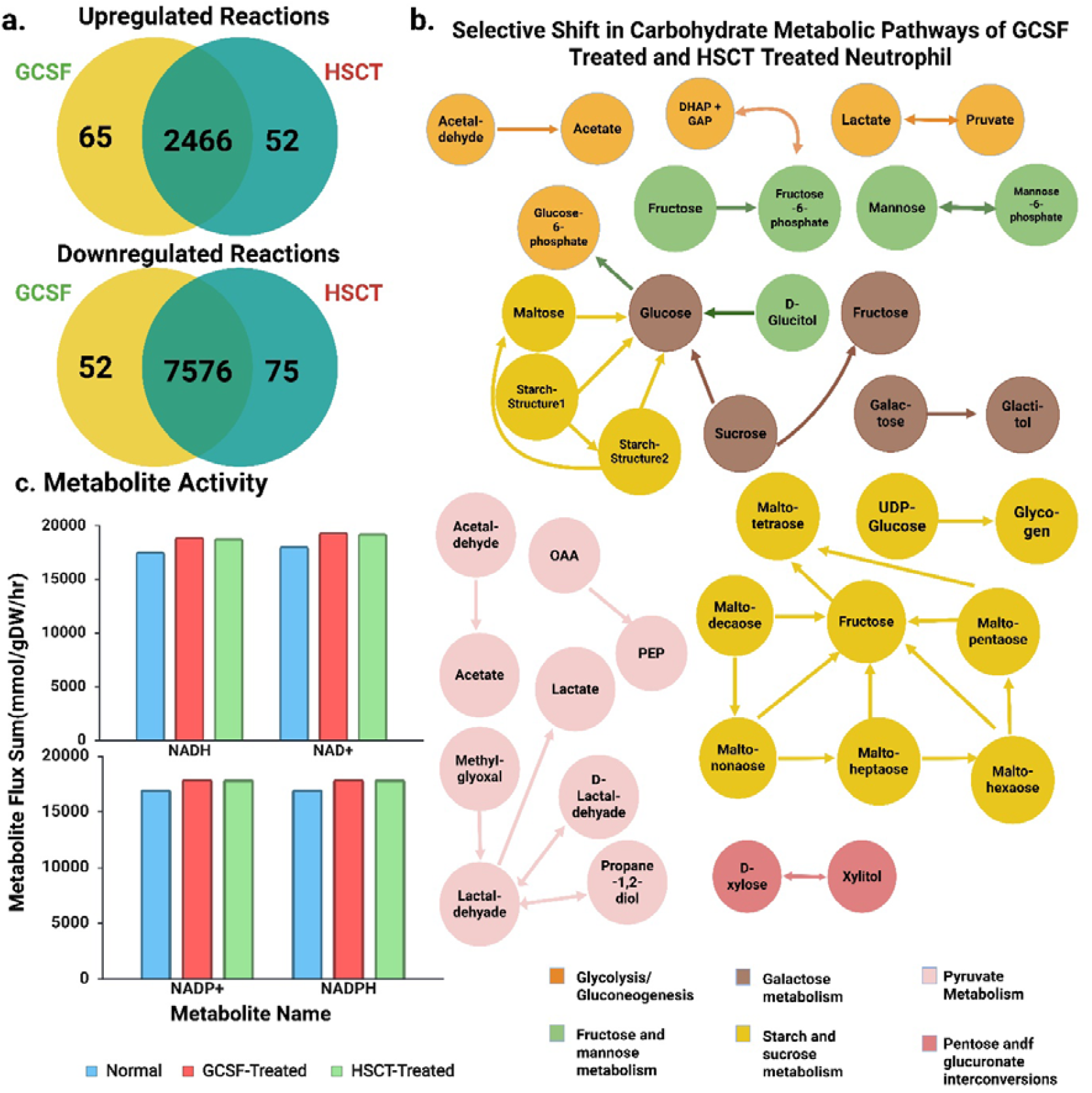
Metabolic flux analysis reveals similarities and differences between G-CSF– treated and HSCT-treated neutrophils. Flux analyses were performed using flux ranges derived from Flux Variability Analysis (FVA). To assess relative changes in flux space, the normal condition was used as the reference baseline. (a) A substantial overlap was observed in both increased and decreased flux reactions between G-CSF–treated and HSCT-treated neutrophils, with only a small subset of reactions unique to each condition. (b) Selective upregulation of carbohydrate metabolism–related reactions suggest adaptive metabolic mechanisms employed by neutrophils in response to distinct stress conditions. Each arrow indicates direction of a reaction, while different metabolic pathways are represented by distinct color. (c) The selective upregulation in carbohydrate metabolism consisted of reactions mainly associated with NADH/NAD+/NADPH/NADP+, which direct impact on the pool sizes of these metabolites.

Similarly, comparison of the flux ranges between the HSCT-treated neutrophil model and the normal condition revealed substantial overlap with the metabolic patterns observed in the G-CSF–treated neutrophil model with minor differences (**Figure 2a**). Additionally, comparative analysis of neutrophils under normal and stimulated conditions revealed a marked increase in metabolic activity centered on carbohydrate utilization and central carbon metabolism. Stimulated neutrophils showed enhanced reactions within glycolysis and gluconeogenesis, including increased interconversion of glycolytic intermediates and elevated flux toward pyruvate-derived end products. Concomitantly, reactions associated with pyruvate metabolism were upregulated, with increased conversion of pyruvate and acetaldehyde to lactate and acetate, indicating a shift toward fermentative or overflow metabolism (**Figure 2b**).

In addition to core glycolytic pathways, stimulated neutrophils exhibited increased activity in starch and sucrose metabolism as well as fructose, mannose, and galactose metabolism, reflecting broader carbohydrate processing capacity. Stepwise degradation of complex carbohydrates and disaccharides to monosaccharides was prominent, suggesting increased mobilization and utilization of available sugar sources. Pentose and glucuronate interconversion pathways were also enriched, consistent with enhanced metabolic flexibility and carbon shunting between pathways. Redox-associated reactions involving NADH/NAD □ and NADPH/NADP□were significantly increased in stimulated neutrophils as shown in Figure 2c. Multiple reactions generating or consuming NADPH in the carbohydrate metabolism were upregulated, indicating elevated redox turnover. The list of reactions upregulated from the carbohydrate metabolism is available in the **Supplementary File**. Together, these findings demonstrate that stimulated neutrophils adopt a highly glycolytic, redox-active metabolic state characterized by rapid energy generation and increased reducing power, consistent with a primed functional phenotype.

The key distinction between the two treated conditions emerged in vitamin metabolism. Vitamin metabolism including Vitamin A, B, C, D & E plays a crucial role in neutrophil functions such as ROS production, migration, proliferation, phagocytosis, and maintaining mitochondrial function. Hence, reprogramming these metabolic pathways could be central to the overall neutrophil metabolism. Both conditions showed elevated fluxes in Vitamin D metabolism, with no changes in Vitamin C metabolism (**Figure 3 a &b**). We observed enhanced fluxes in Vitamin A, and Vitamin E but reduced fluxes for Vitamin B for G-CSF treated neutrophils. While in HSCT-treated neutrophils, Vitamin E metabolism now showed reduced flux space and Vitamin A remained same to the normal condition, while the activity of other vitamin metabolism remained similar to that of G-CSF treated neutrophils as shown in **Figure 3 a & b**.

**Figure 3:**
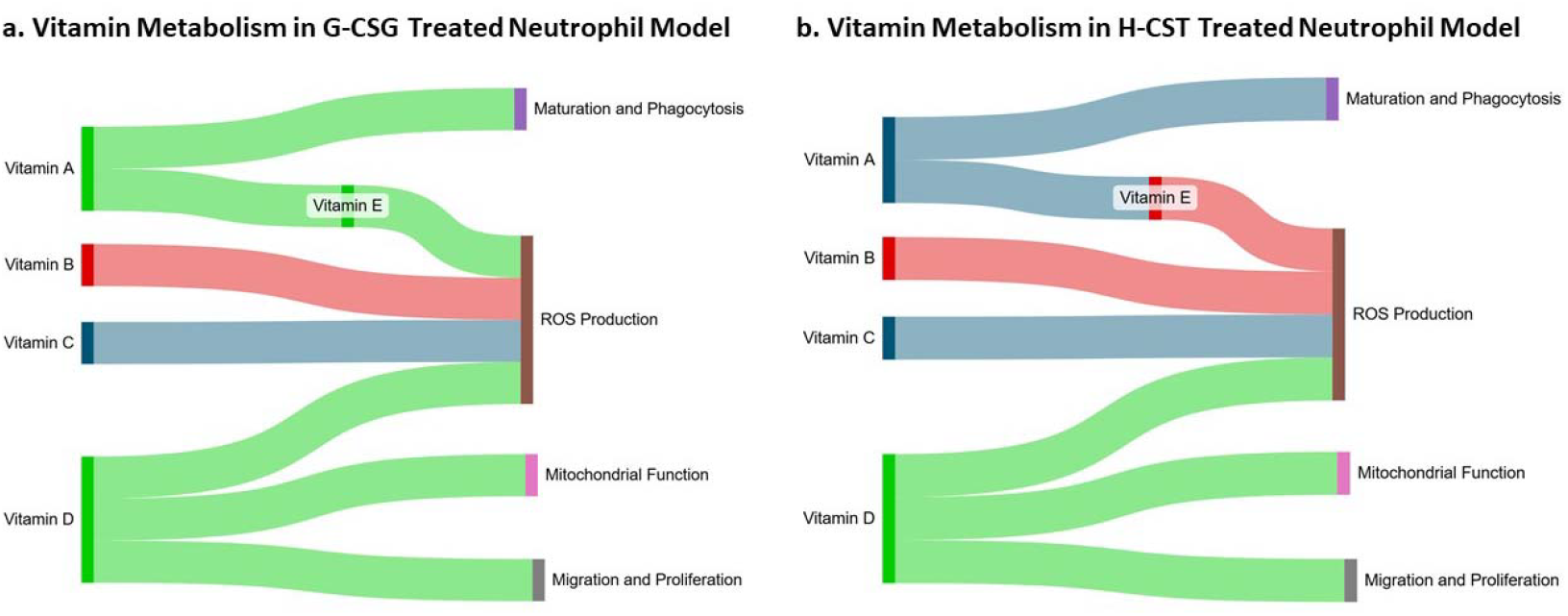
Alterations in vitamin metabolism in G-CSF– and HSCT-treated neutrophils and their potential impact on overall neutrophil metabolism and overall immune function. (a) Increased or decreased overall fluxes in the different vitamin metabolism in the G-CSF Treated neutrophil model and their connection key neutrophil functions. (b) Increased or decreased overall fluxes in the different Vitamin metabolism in the HSCT Treated neutrophil model and their connection key neutrophil functions. Together, these findings indicate that neutrophils adopt condition-specific metabolic strategies to accommodate stress-induced metabolic shifts. The metabolism highlighted by using green shows increased flux activity in the pathway and red denote decreased flux activity. Blue shows the flux space that did not change in comparison to the normal condition.

Additionally, we also observed distinct differences in exchange and transport reactions. G-CSF–treated neutrophils exhibited expanded flux spaces for the exchange of compounds such as gamma-tocopherol, gamma-tocotrienol, lipoic acid, quinolinate, 3,4-dihydroxymandelate, and 5-hydroxytryptophol. In contrast, these reactions displayed reduced flux ranges in the HSCT-treated neutrophil model. Instead, HSCT-treated neutrophils showed increased activity in the exchange or transport of retinol, aquacob(III)alamin, 5-hydroxyindoleacetate, 2-methylbutyrylglycine, isobutyrylglycine, glucuronide–thiomethyl-acetaminophen conjugate, thiomethyl-sulphoxide–acetaminophen–glucuronide, thiomethyl-sulphoxide–acetaminophen– sulphate, and retinyl palmitate. These key metabolites impact pathways such as tryptophan metabolism, and Vitamin metabolism. These observations delineate that despite having substantial overlap in metabolic shift when comparing G-CSF and HSCT treated neutrophils to normal conditions, they also have their unique metabolic properties. This is mostly evident through each model’s preference for specific metabolites as shown by transport and exchange reaction fluxes.

### Metabolic Reprogramming of Neutrophil in Pancreatic Cancer

In PDAC neutrophil model, the observed metabolic shift is very different compared to G-CSF-treated, and HSCT-treated condition as shown in **Supplementary Figure 1a & b**. Over 80% of the total reactions showed widened flux space, including the full glycolysis/gluconeogenesis, TCA cycle, and the majority of amino acid metabolism. Very small number of reactions show shrunk flux Space (only around 560, sparsely distributed among all pathways). In pancreatic cancer (PDAC), overall higher flux compared to normal condition indicating a profound, systemic metabolic hyperactivation in tumor-associated neutrophils (TANs). This scenario aligns with observed reprogramming in the tumor microenvironment (TME), where TANs adapt to hypoxia, nutrient competition, and cytokine signals to support pro-tumor functions. While no studies explicitly quantify exactly 80% of reactions with elevated fluxes in PC neutrophils, inferential evidence from single-cell omics, metabolomics, and metabolic modeling supports that a majority of pathways including glycolysis, OXPHOS, PPP, and amino acid catabolism are upregulated, often by several-fold, to meet heightened demands in cancer conditions (**Supplementary Figure 1b)**. The major difference we noted between other condition neutrophils and PDAC neutrophil is the increased flux of selective reactions from a pathway vs full pathway upregulated in PDAC models.

On the other hand, reactions that showed shrunk flux spaces are mostly exchange/demand, and transport reactions. Such reactions were associated with important compounds such as D-arginine, ornithine, eumelanin, maltodecaose, maltotriose, umbelliferone, corticosterone, aldosterone, retinol, VLDL, chylomicron, and so on as shown in Supplementary **Figure 1c**. Specially, reduced fluxes of crucial compounds such as arginine, and ornithine exchange could indicate enhanced local arginine depletion, promoting immunosuppression and tumor progression. Additionally, compounds such as VLDL, coumarin, cholesterol, retinol, and glycogenin G4G7 also play important roles in modulating several key functions of neutrophils as shown in Supplementary Figure 1c. Together, the reduced fluxes across these diverse compounds suggest a “deprived” or “minimalist” state in TANs, where neutrophils adapt to TME constraints by downregulating non-essential exchanges, focusing on core survival pathways such as central carbon metabolism, amino acid metabolism, and fatty acid metabolism. This mirrors observations in PDAC, where TAN subpopulations show hyper-glycolysis and overall hyper-activated state. Additionally, our observations hint towards immunosuppression possibly due to arginine and, ornithine exchange shifts that directly impact Nitric Oxide (NO) cycle. The details of the flux comparison are available in **Supplementary File**.

Hence, based on our flux analysis we provide additional support that neutrophils metabolically reprogram themselves to aid tumor, as evidenced by the “revved up” metabolic pathways while reducing their dependency in the harsh external environment as seen in the reduced flux space of exchange/demand reactions of key metabolites.

### Solute Carrier Transporter Family at the Heart of Neutrophil Metabolic Reprogramming

We examined metabolic reprogramming in neutrophils under G-CSF treatment, HSCT, and pancreatic ductal adenocarcinoma (PDAC) conditions. While neutrophils from G-CSF– and HSCT-treated samples exhibited notable similarities, their overall metabolic profiles remained distinct from those observed in PDAC, where neutrophils displayed the most divergent metabolic behavior. Notably, reactions that were upregulated in G-CSF and HSCT conditions showed a contraction of flux space in PDAC, and vice versa. In addition, certain reactions exhibited expanded flux space in one condition while being constrained in the other two relative to the normal state. These observations prompted us to investigate whether common underlying factors could explain the global metabolic differences across conditions.

Our analysis revealed that the transport of several key metabolites—including fructose, glucose, noradrenaline, serotonin, acetylcholine, prostaglandin E2, prostaglandin F2α, dopamine, glutamate, aspartate, histamine, and multiple fatty acids—displayed distinct condition-specific behavior. Importantly, many of these transport reactions were mediated by Solute Carrier (SLC) family transporters, particularly members of the SLC0, SLC1, SLC2, SLC8, SLC22, SLC25, SLC26, and SLC27 families. Transporters such as SLC1A1, SLC1A2, SLC1A3, and SLC1A6 regulate glutamate and neutral amino acid transport, which are essential for cellular energy production and antioxidant defense. Additionally, SLCO2A1 mediates prostaglandin E2 transport, a critical regulator of neutrophil immune function, especially in inflammatory and cancer contexts. The transport of other key metabolites, including carnitine, glutamate, and sodium, was also predominantly governed by SLC family members.

We further investigated which metabolite classes were most affected by these shifts in exchange of reaction fluxes. Given that SLC transporter genes are known to function cooperatively rather than independently, as illustrated in **Figure 4a**, we observed pronounced differences in the abundance of amino acids such as glutamate, aspartate, and L-carnitine, as well as several fatty acids, including (10Z)-heptadecenoic acid, 9-heptadecylenic acid, mead acid, and cis-cetoleic acid (**Figure 4b**). Altered transport of these metabolites likely plays a critical role in shaping the overall metabolic phenotype of neutrophils.

**Figure 4:**
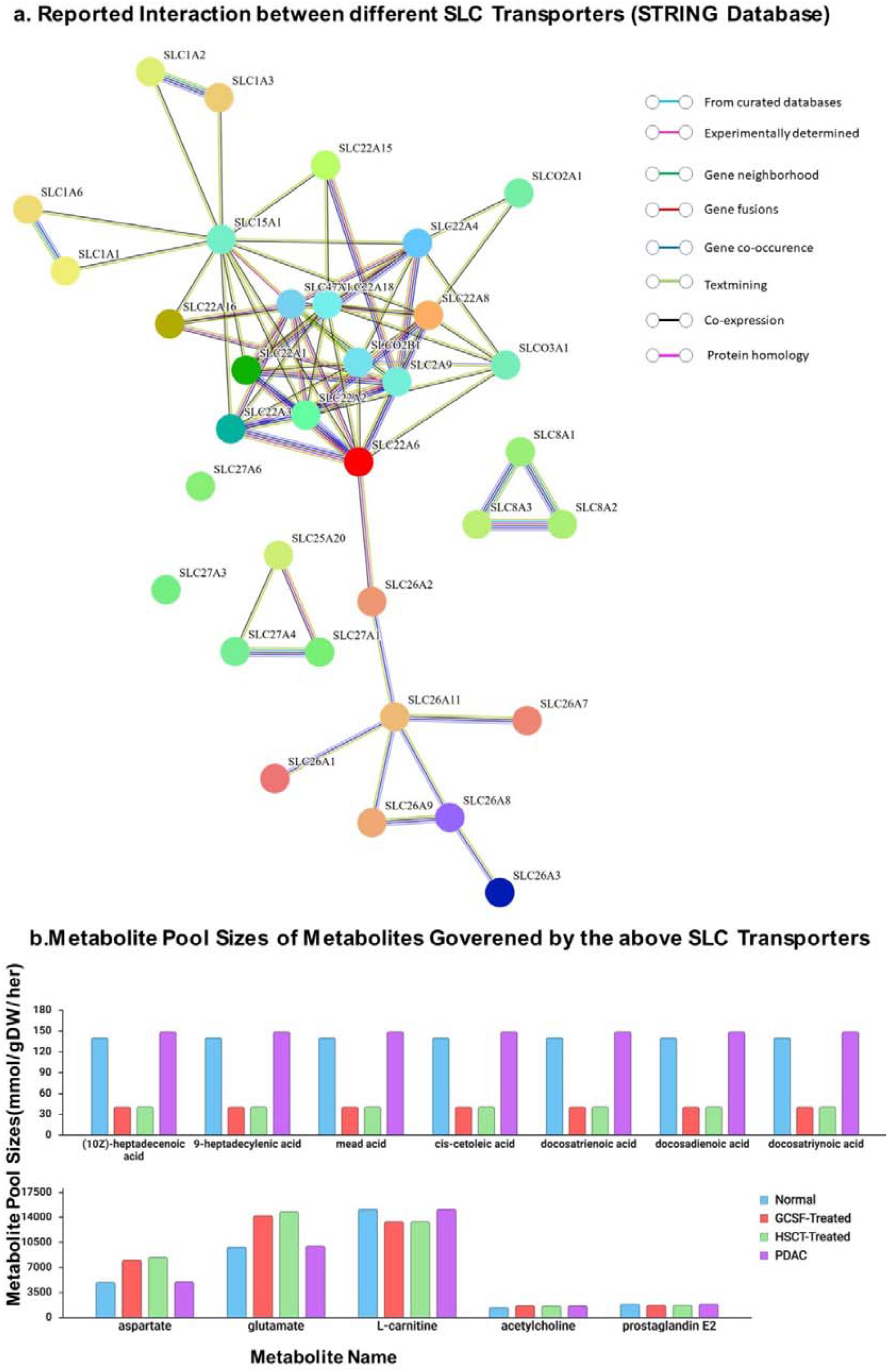
Influence of the Solute Carrier Family Transporters. (a) Reported Interaction between the different genes identified by the flux analysis from SLC Family from the STRING database. (b) Most profound changes in Metabolite Pool size of compounds that are governed by the SLC family transporters. The metabolite pool sizes are obtained from Flux Sum Analysis. Most drastic changes can be seen in fatty acids and key amino acids such as aspartate, and glutamate with notable changes in key metabolites such as carnitine, prostaglandin, and acetylcholine.

While alterations in central carbon metabolism under stress conditions are well documented, our findings indicate that vitamin metabolism also plays a significant role in neutrophils following G-CSF and HSCT treatments. Consistent with previous studies, prostaglandin E2 emerged as an important regulator of neutrophil function. Integrating these observations, we propose that SLC transporter activity may complement and fine-tune neutrophil metabolic capacity across conditions. In G-CSF– and HSCT-treated neutrophils, the majority of pathways exhibited reduced flux space, indicative of diminished overall metabolic activity; however, selective upregulation of carbohydrate metabolism supports elevated reactive oxygen species (ROS) production, while shifts in vitamin metabolism appear to mitigate ROS-induced stress. Moreover, flux analysis suggests that neutrophils under these conditions rely more heavily on external nutrient availability, as reflected by increased exchange reaction activity (Figure 5 a).

**Figure 5:**
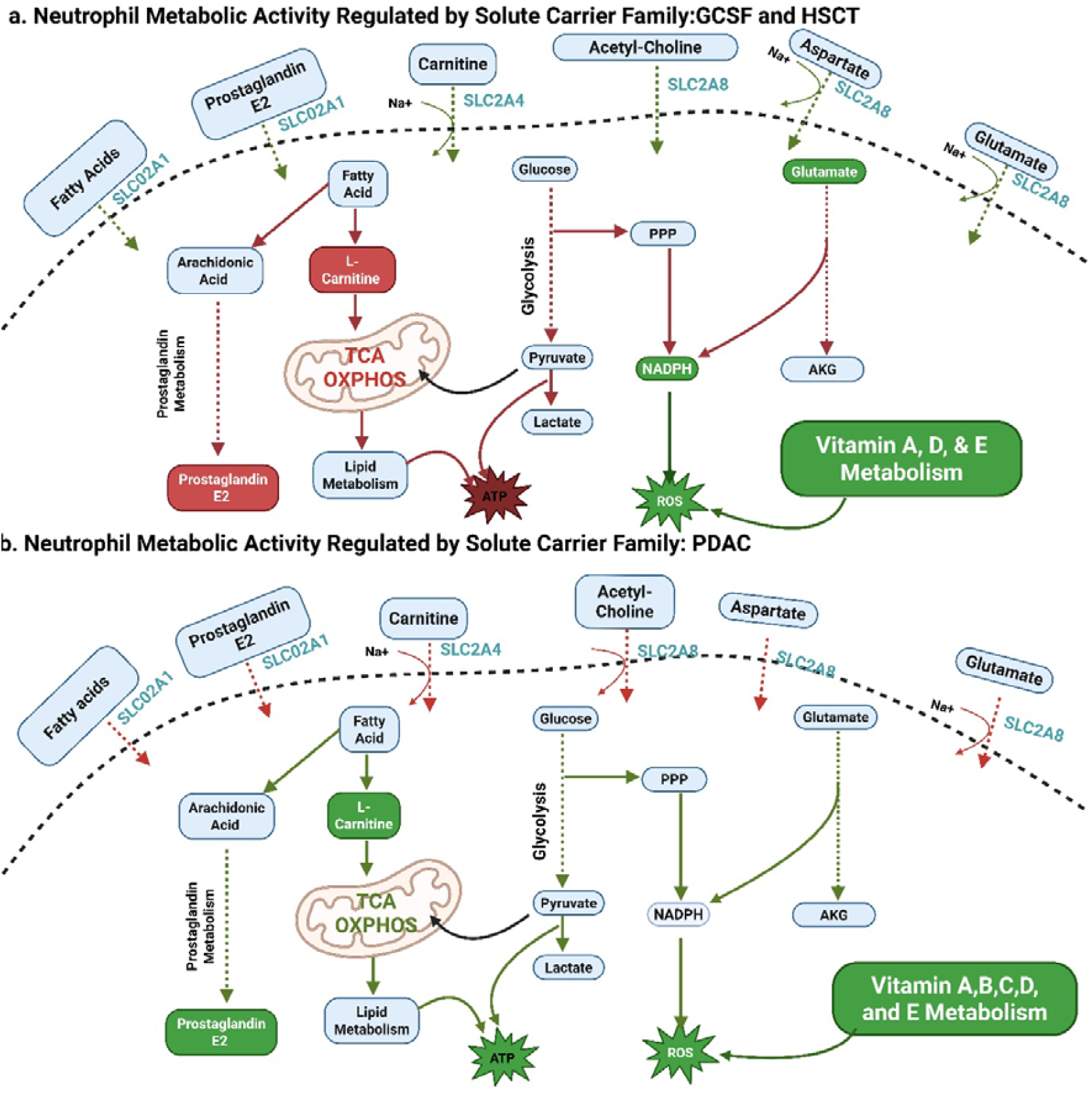
Schematic of the impact of the shift in transport of compounds governed by SLC Family transporters. (a) The increased flux activity in the transport of the metabolites aids the diminished metabolic activity of the neutrophils after G-CSF and HSCT treatment.(b)The reduced flux of the transporters here along with most of the metabolic pathways suggests a reduced dependency on the external environment and contributes to the immunosuppression during PDAC progression.

In contrast, neutrophils in the PDAC microenvironment appear to upregulate intrinsic metabolic pathways while limiting dependence on extracellular resources, likely as an adaptive response to harsh tumor conditions (**Figure 5b**). Across all contexts, SLC transporters seems to play a central role by regulating the transport of critical metabolites including glutamate, aspartate, fatty acids and key ions such as sodium and potassium, thereby orchestrating condition-specific neutrophil metabolic reprogramming. To further assess whether SLC transporters can influence metabolic function, we constrained the flux of SLC transporter–mediated reactions (26 reactions, including transport/exchange of glycogenin G4G7, mead acid, aspartate, glutamate, prostaglandins, among others) in G-CSF and HSCT models to match the flux ranges observed in PDAC models (the details are available in Supplementary File). Under these constraints, we observed substantial shifts in the flux space of multiple pathways, including starch and sucrose metabolism, purine metabolism, glycerophospholipid metabolism, retinol metabolism, and vitamin metabolism in G-CSF and HSCT neutrophils. These findings indicate that SLC transporters play a significant role in regulating overall metabolic activity.

## Discussion

Neutrophils exhibit remarkable metabolic plasticity, dynamically rewiring their bioenergetic and biosynthetic programs to support diverse functional states, including granulopoiesis, antimicrobial defense, and tumor-associated phenotypes^9^. Recent studies have highlighted neutrophil plasticity driven by signaling and transcriptional reprogramming; however, the specific scope and contribution of metabolic regulation have not been systematically characterized^8^. In this study, we provide a comprehensive analysis of neutrophil metabolic remodeling across distinct stress conditions: G-CSF treatment, HSCT treatment, and pancreatic ductal adenocarcinoma (PDAC). Using flux variability analysis (FVA) ranges and comparisons to a normal baseline, we identify condition-specific alterations that reveal how metabolic networks adapt to activation, recovery, or tumor-imposed constraints.

G-CSF and HSCT treatments are used to restore neutrophil numbers and immune competence in conditions where hematopoiesis is compromised^13,17^. Despite their clinical benefits, both treatments have important limitations^34,35^. G-CSF-induced neutrophils often exhibit altered functional and metabolic states, which may compromise antimicrobial efficiency or promote excessive inflammation under certain conditions^34^. HSCT-associated neutrophils arise in a highly stressed, inflammatory, and nutrient-restricted environment, frequently leading to immature or dysregulated phenotypes and increased susceptibility to infection or inflammatory complications^14^. Hence, by using systems biology approaches such as genome-scale metabolic (GSM) modeling we can exhaustively analyze the metabolic capabilities of neutrophils after each treatment, highlight possible similarities as well as differences and uncover possible strategies to mitigate the harmful effects^19^. However, it is extremely important to acknowledge the inherent limitations of GSM models. Often their predictive power is limited by simplified assumptions, such as steady-state fluxes, incomplete pathway coverage, and lack of enzyme kinetics or dynamic regulation^36^. Model predictions are also highly sensitive to environmental conditions and often cannot fully capture context-specific stress responses or microenvironmental influences^37^. Despite these limitations, GSMs remain valuable for identifying key metabolic shifts and guiding experimental hypotheses^38^. To this end, our flux analysis identifies the key metabolic vulnerabilities of neutrophils in each condition and highlights the role of Vitamin metabolism as the main distinction between the two conditions.

In G-CSF–treated neutrophils, increased fluxes involving gamma-tocopherol and gamma-tocotrienol, both vitamin E isoforms with potent antioxidant properties suggest an adaptive response to elevated reactive oxygen species (ROS) production^39^. G-CSF is known to prime neutrophils for enhanced antimicrobial activity, including increased superoxide generation, which imposes substantial oxidative stress^12^. Upregulation of vitamin E–associated metabolic pathways may therefore serve a protective role by scavenging ROS and stabilizing cellular membranes^39^. Similarly, enhanced lipoic acid metabolism in G-CSF–treated neutrophils likely support mitochondrial function and redox balance, consistent with the dual role of lipoic acid as both an antioxidant and a mitochondrial enzyme cofactor^40^. In addition, G-CSF–driven emergency granulopoiesis commonly occurs in inflammatory settings that activate the kynurenine pathway of tryptophan metabolism.^41^ The observed increase in quinolinate-associated fluxes in G-CSF–treated neutrophils is therefore consistent with heightened immune activation, as quinolinate functions as an immunomodulatory metabolite and may contribute to neutrophil functional reprogramming^42^. Although G-CSF–treated and HSCT-treated neutrophils share several metabolic features, key differences emerge that reflect their divergent functional states. In G-CSF–treated neutrophils, increased fluxes involving gamma-tocopherol and gamma-tocotrienol both vitamin E isoforms with potent antioxidant properties suggest an adaptive response to elevated reactive oxygen species (ROS) production^12^. G-CSF is known to prime neutrophils for enhanced antimicrobial activity, including increased superoxide generation, which imposes substantial oxidative stress^43^.

In contrast, neutrophils following HSCT exhibit a metabolic profile consistent with impaired activation capacity and a shift toward survival and recovery^35^. HSCT conditioning regimens induce profound oxidative stress and ROS-mediated tissue damage, requiring neutrophils to counteract oxidative burden during early engraftment^15^. This process can rapidly deplete endogenous antioxidant reserves, resulting in reduced flux through antioxidant-related pathways, including uptake, recycling, and exchange reactions captured in our model (e.g., MAR09153, MAR09152, MAR09167)^17^. Such reductions likely reflect both antioxidant exhaustion and metabolic reprioritization away from activation-associated processes^15^. Alterations in tryptophan metabolism further distinguish post-HSCT neutrophils from their G-CSF–treated counterparts^41^. Reduced flux through quinolinate-producing pathways may arise from suppression of upstream kynurenine metabolism, a phenomenon reported in HSCT patients, and linked to systemic inflammation, immune suppression, and microbiome disruption^41^. These factors, particularly gut dysbiosis affecting indole metabolism, are common after HSCT and may contribute to diminished amino acid catabolism and attenuated inflammatory signaling in reconstituting neutrophils^35^. Additional metabolic contractions observed post-HSCT involve catecholamine and serotonin metabolism. Reduced fluxes through catecholamine degradation intermediates, such as 3,4-dihydroxymandelate, may reflect autonomic dysfunction, diminished sympathetic signaling, or the effects of immunosuppressive therapies (e.g., calcineurin inhibitors), all of which are known to impair neutrophil function^44,45^. Likewise, decreased flux through 5-hydroxytryptophol may result from redirection of serotonin metabolism or reduced gut-derived serotonin production, consistent with antibiotic exposure, graft-versus-host disease, and microbiome perturbations commonly observed following HSCT^46^.

On the other hand, the almost entirely enhanced flux space in PDAC neutrophil models indicates a hallmark of an increased metabolic demand in the TME environment^47^. The amplified fluxes supporting ROS generation support neutrophil extracellular traps (NETs) production and immunosuppression^48^. In nutrient-scarce areas, fluxes shift toward glutamine-fueled OXPHOS for efficient ATP, with antioxidants (e.g., glutathione via PPP) upregulated to counter ROS overload^49,50^. In PDAC, particularly fatty acid oxidation (cytosolic), carnitine shuttle, as well as the role of SLC transporter family, was found to be a very important factor in progression of cancer itself^25^. However, GSM models are not without their own limitations. Most GSM models assumes a steady-state system and therefore is uncapable of capturing dynamic metabolic changes over time. They also lack detailed regulatory mechanisms, such as gene expression control and signaling pathways, which can lead to biologically unrealistic predictions. Additionally, GSMs depend on the completeness and accuracy of existing metabolic reconstructions, which are often incomplete, especially for less-studied cell types, resulting in missing or incorrect reactions. Furthermore, they often oversimplify cellular compartmentalization and struggle to fully incorporate cell-type specificity and heterogeneous experimental data, making careful interpretation and experimental validation essential. Despite their limitations, GSM models remain highly valuable because they provide a systematic and predictive framework for analyzing metabolism at a scale that is difficult to achieve experimentally. They are especially useful for integrating large-scale omics data to uncover metabolic rewiring, compare cell states, and prioritize key enzymes or pathways for experimental validation. Even when not perfectly accurate, they serve as powerful hypothesis-generating tools that guide experiments, reduce search space, and accelerate discovery, particularly in complex systems like immune cell metabolism.

Given these advantages, genome-scale metabolic models (GEMs) are well-suited for studying neutrophils, which are highly metabolically active cells that depend on solute carrier transporters for nutrient and ion uptake. These processes support rapid energy production through pathways such as glycolysis and fatty acid oxidation, as well as essential effector functions including phagocytosis, NETosis (neutrophil extracellular trap formation), and migration^51^. G-CSF is a key cytokine that mobilizes neutrophils from bone marrow, enhances survival, and primes them for inflammation often used in HSCT to accelerate engraftment and reduce neutropenia^14^. In contrast, in pancreatic ductal adenocarcinoma (PDAC), tumor-associated neutrophils (TANs) often adopt a pro-tumor, immunosuppressive phenotype, with altered metabolism that could involve change in SLC activity^25,52,53^. Recent advancements have shed lights into the importance of SLC family and their role in shaping the metabolism of immune cells such as macrophages, T cells, B cells, etc but their specific role in neutrophil metabolism still remains elusive^52,54^. To this end, we found that the SLC transporter family actively shapes the metabolic function of neutrophil in different conditions.

Our findings suggest divergent regulatory programs involving SLC transporters across conditions. In G-CSF treatment and HSCT-treatment, upregulation of SLC-mediated fluxes could be a result of cytokine signaling, particularly STAT3/JAK pathways, to enhance nutrient and ion transport required for inflammation and recovery^8,55^. In contrast, PDAC-associated neutrophils display inhibition of multiple SLC pathways, potentially due to tumor hypoxia, stromal interactions, or epigenetic repression^25^. This suppression may promote a shift toward oxidative phosphorylation and pro-tumor neutrophil phenotypes, providing a mechanistic explanation for the adverse outcomes observed with G-CSF administration in neoadjuvant PDAC settings^31,56^. Notably, SLC transporters such as SLCO2A1 (PGT), SLC22A8 (OAT3), and SLC22A4 (OCTN1) may form a functionally relevant network in neutrophils^55^. Although direct physical interactions have not been established, these transporters share overlapping substrates, inflammatory regulation, and expression patterns in immune cells. In tumor-associated neutrophils, reduced SLCO2A1 activity could impair prostaglandin E□ (PGE□)clearance, reinforcing immunosuppressive signaling within the tumor microenvironment^57,58^. Collectively, our findings highlight how neutrophil metabolism is finely tuned to context-specific demands and suggest that metabolic flux patterns may serve as indicators of immune competence, recovery status, or pathological reprogramming across diverse clinical settings.

## Methods

### Genome-scale Metabolic Models Generation

Single-cell RNA-sequencing (scRNA-seq) data for neutrophils under four conditions: steady state (Normal), granulocyte colony-stimulating factor (G-CSF) treatment, hematopoietic stem cell transplantation (HSCT), and pancreatic ductal adenocarcinoma (PDAC) were obtained from Machado et al. (2022)^8^. The dataset comprised neutrophils isolated from peripheral blood and bone marrow samples across steady-state and stress conditions.

Condition-specific neutrophil datasets were generated using the Seurat pipeline in RStudio^59^. Cells with fewer than 5% mitochondrial transcripts were excluded, and gene expression was normalized using the LogNormalize method in Seurat^59^. The neutrophils were identified based on the markers presented by Machado et al. (2022)^8^. These datasets were subsequently used to reconstruct four context-specific neutrophil metabolic models using the ftINIT pipeline^27^, with the Human1^26^ genome-scale metabolic model serving as the reference network. The ftINIT pipeline was implemented using the RAVEN toolbox with default parameters unless otherwise specified. Optimization problems were solved using the Gurobi solver^27^.

Additionally, Gene–protein–reaction (GPR) associations and condition-specific metabolic gene expression (average gene expression for each condition) derived from the scRNA-seq data were used to define reaction-specific upper and lower flux bounds^60^. Reaction flux bounds were scaled proportionally to gene expression levels using GPR rules, such that higher expression corresponded to relaxed flux constraints, while reactions associated with lowly expressed genes were restricted. The updated upper and lower bounds of each condition are available in **Supplementary File**.

All reconstructed models employed a common biomass objective function for immune cells, originally described by Bordbar et al. (2010)^61^. The final context-specific models each contained over 11,000 reactions and more than 7,000 metabolites. Comprehensive quality control and curation were performed to ensure network connectivity and the inclusion of biologically relevant pathways, reactions, and metabolites. Complete lists of genes, reactions, and metabolites for each model are provided in the Supplementary File. All reconstructed models are provided in SBML format, and the code used for preprocessing and model reconstruction is available at https://github.com/ssbio/Neutrophil.

### Flux balance, flux sum, flux variability analysis

Flux Balance Analysis (FBA)^29^ is employed in this study to examine metabolite flow under different conditions. FBA is a widely used computational approach for analyzing biochemical networks, particularly genome-scale metabolic (GSM) models, which encompass known metabolic reactions in a biological system along with their associated genes and enzymes^29^. The GSM model is represented by a stoichiometric matrix, with metabolites and reactions defining its structure, while upper and lower bounds are applied to each reaction to impose constraints based on nutrient availability and other microenvironmental or genetic conditions. FBA computes a flux value for every reaction in the network^62^.

Flux Variability Analysis (FVA)^28^, an extension of FBA, determines the minimum and maximum feasible fluxes for each reaction under a given condition. In addition, Flux Sum Analysis (**FSA**) is used to estimate metabolite pool sizes across different conditions^63^. All the scripts are available at https://github.com/ssbio/Neutrophil.

## Data Availability

All the generated and used data information is provided in this paper and its Supplementary Files. Additionally, the GSM models (xml versions), and the flux analysis results can be found included in the manuscript, Supplementary files or in this GitHub repository: https://github.com/ssbio/Neutrophil.

## Code Availability

All the codes used in this study are available at https://github.com/ssbio/Neutrophil.

## Acknowledgements

We gratefully acknowledge the funding support from the National Institute of Health (NIH) R35 MIRA grant (5R35GM143009), awarded to RS. We would also like to thank the Holland Computing Center (HCC) of the University of Nebraska, which receives support from the Nebraska Research Initiative (United States of America).

## Author Contributions

S.M. worked on concept development for this work. The analysis followed by the writing was done by S.M. R.S provided meaningful insights in the writing, editing, and model analysis process.

